# Genetic background and diet affect brown adipose gene co-expression – metabolic associations

**DOI:** 10.1101/2019.12.20.884197

**Authors:** Caryn Carson, Heather A Lawson

## Abstract

Adipose is a dynamic endocrine organ that is critical for regulating metabolism and is highly responsive to nutritional environment. Brown adipose tissue is an exciting potential therapeutic target, however there are no systematic studies of gene-by-environment interactions affecting function of this organ. We leveraged a weighted gene co-expression network analysis to identify transcriptional networks in brown adipose tissue from LG/J and SM/J inbred mice fed high or low fat diets, and correlate these networks with metabolic phenotypes. We identified 8 primary gene network modules associated with variation in obesity and diabetes-related traits. Four modules were enriched for metabolically relevant processes such as immune and cytokine response, cell division, peroxisome functions, and organic molecule metabolic processes. The relative expression of genes in these modules is highly dependent on both genetic background and dietary environment. Genes in the immune/cytkine response and cell division modules are particularly highly expressed in high fat-fed SM/J mice, which show unique brown adipose-dependent remission of diabetes. The interconnectivity of genes in these modules is also heavily dependent on diet and strain, with most genes showing both higher expression and co-expression under the same context. We highlight 4 candidate genes, *Col28a1, Cyp26b1, Bmp8b*, and *Kcnj14*, that have distinct expression patterns among strain-by-diet contexts and fall under metabolic QTL previously mapped in an F_16_ generation of an advanced intercross between these two strains. Each of these genes have some connection to obesity and diabetes-related traits, but have not been studied in brown adipose tissue. In summary, our results provide important insights into the relationship between brown adipose and systemic metabolism by being the first gene-by-environment study of brown adipose transcriptional networks and introducing novel candidate genes for follow-up studies of biological mechanisms of action.

**Author Summary:** Research on brown adipose tissue is a promising new avenue for understanding and potentially treating metabolic dysfunction. However, we do not know how genetic background interacts with dietary environment to affect the brown adipose transcriptional response, and how this might affect systemic metabolism. Here we report the first investigation of gene-by-environment interactions on brown adipose gene expression networks associating with multiple obesity and diabetes-related traits. We identified 8 primary networks correlated with variation in these traits in mice, including networks enriched for immune and cytokine response, cell division, organic molecule metabolism, and peroxisome genes. Characterizing these networks and their distinct diet-by-strain expression and co-expression patterns is an important step towards understanding how brown adipose tissue responds to an obesogenic diet, how this response affects metabolism, and how this can be modified by genetic variation.

## Introduction

Obesity and associated metabolic disorders are reaching epidemic prevalence world-wide. While some of this prevalence is the result of increasingly inactive lifestyles and changing dietary norms, the hundreds of genome-wide association study (GWAS) ‘hits’ for obesity and metabolic diseases indicate there is also a strong genetic component [1,2]. Thus it is critical to understand how gene-by-environment interactions are contributing to metabolic dysfunction. Such interactions have been shown to underlie variation in obesity and diabetes risk [3–8], and many individual genes have been identified with natural variants in human populations affecting metabolic response to environmental perturbations [9–14]. Research in animal models, in particular mouse models, has been used to manipulate dietary intake and environment in order to better understand the gene-by-environment interactions most relevant to human metabolism [15–19]. Frequently, studies of obesity in mice look at adipose tissue as a primary metabolic organ, with relatively recent focus on brown adipose tissue as a pro-health therapeutic target for obesity [20].

Brown adipose tissue is distinct from white adipose tissue and has mostly been studied for its role in non-shivering thermogenesis, the release of energy as heat through the activity of UCP1 [21,22]. Brown adipose tissue is found in adult humans [23,24] and increased brown adipose activity is associated with a healthier metabolic profile [25] and lower body fat percentage [26]. It is also associated with amelioration of elevated plasma lipid levels in a hyperlipidemic mouse model [27] and remission of the metabolic dysfunction associated with impaired pancreatic islet function in a mouse model of type I diabetes [28]. Mice lacking brown adipose tissue develop obesity and metabolic dysfunction [29] that is independent from the loss of thermogenic UCP1 activity [21,30,31], indicating that brown adipose contributes to healthy metabolism through thermogenesis-independent mechanisms. Several studies have sought to identify potential metabolically-relevant brown adipose cytokines, or “batokines” underlying these mechanisms [32–34]. However, very little research has focused on understanding the potential regulation of these batokines, and most studies regarding transcriptional networks in brown adipose tissue have focused exclusively on identifying regulators and effectors of nonshivering thermogenesis and brown adipocyte identity [35,36,45–49,37–44].

We wanted to more broadly understand the transcriptional networks existing within brown adipose tissue, to investigate how these networks are affected by genetic background and dietary environment, and determine how they associate with metabolic variation. To do this, we chose to study SM/J and LG/J inbred mice fed high and low fat diets. These strains both respond to a high fat diet with obesity, elevated fasting glucose, and impaired glucose tolerance at 20 weeks of age [50]. However, by 30 weeks, high fat-fed SM/J mice resolve their glycemic dysfunction [51]. This occurs concurrently with a dramatic expansion of their interscapular brown adipose depots, making SM/J mice a unique and intriguing model system in which to investigate brown adipose transcriptional networks, how they correlate with metabolic traits, and how they are affected by dietary environment. In this study, we employed a weighted gene co-expression network analysis (WGCNA) [52,53] and identified eight primary gene modules within brown adipose tissue that correlate with one or more obesity and diabetes-related phenotypes. The expression profile of genes within these modules is dependent on both strain and dietary contexts, indicating gene-by-environment interactions contribute significantly to variation in brown adipose tissue function. This study is an important first step in elucidating metabolically-relevant brown adipose transcriptional networks, how they are affected by genetic background and diet, and how they contribute to systemic metabolism through mechanisms beyond thermogenesis.

## Results

### Brown adipose expression and metabolic traits vary among strain and diet contexts

To understand how brown adipose gene expression in different genetic backgrounds contributes to metabolic variation in different environmental contexts, we raised LG/J and SM/J mice on isocaloric high and low fat diets. The mice were extensively phenotyped (**Supplementary Table 1**) and RNA sequencing of brown adipose was used to assess both mRNA and noncoding RNA transcript levels. To understand how genetic background and diet interact to affect the brown adipose transcriptome, samples were clustered based on gene expression (**Figure 1, top**). Strain is a much better predictor of overall clustering than diet, with the LG/J samples clustering into a single group comprising both diets. The SM/J samples separate into two main clusters, one for each diet, indicating that the SM/J’s brown adipose is more responsive to dietary environment. A heatmap is used to visualize the metabolic variation among the strains and diets (**Figure 1, bottom**). Glucose and insulin parameters consistently show higher values in the high fat-fed SM/J’s relative to low fat-fed mice or LG/J mice on either diet. Lipid levels are more mixed and variable throughout the entire population. LG/J mice have lower brown adipose to body weight ratios than SM/J mice. Within the SM/J’s, high fat-fed mice have both higher body weight and higher brown adipose to body weight ratios than low fat-fed mice.

**Table 1.**
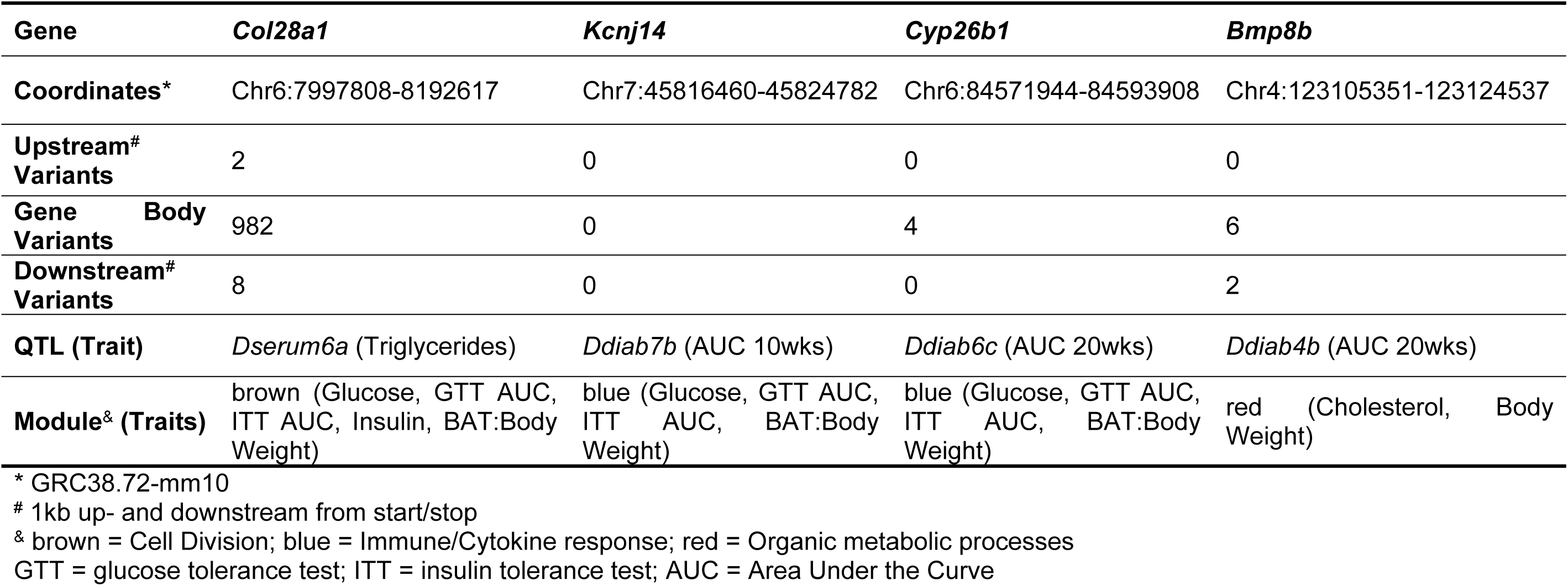
Differentially connected genes falling in metabolic QTL.

**Figure 1:**
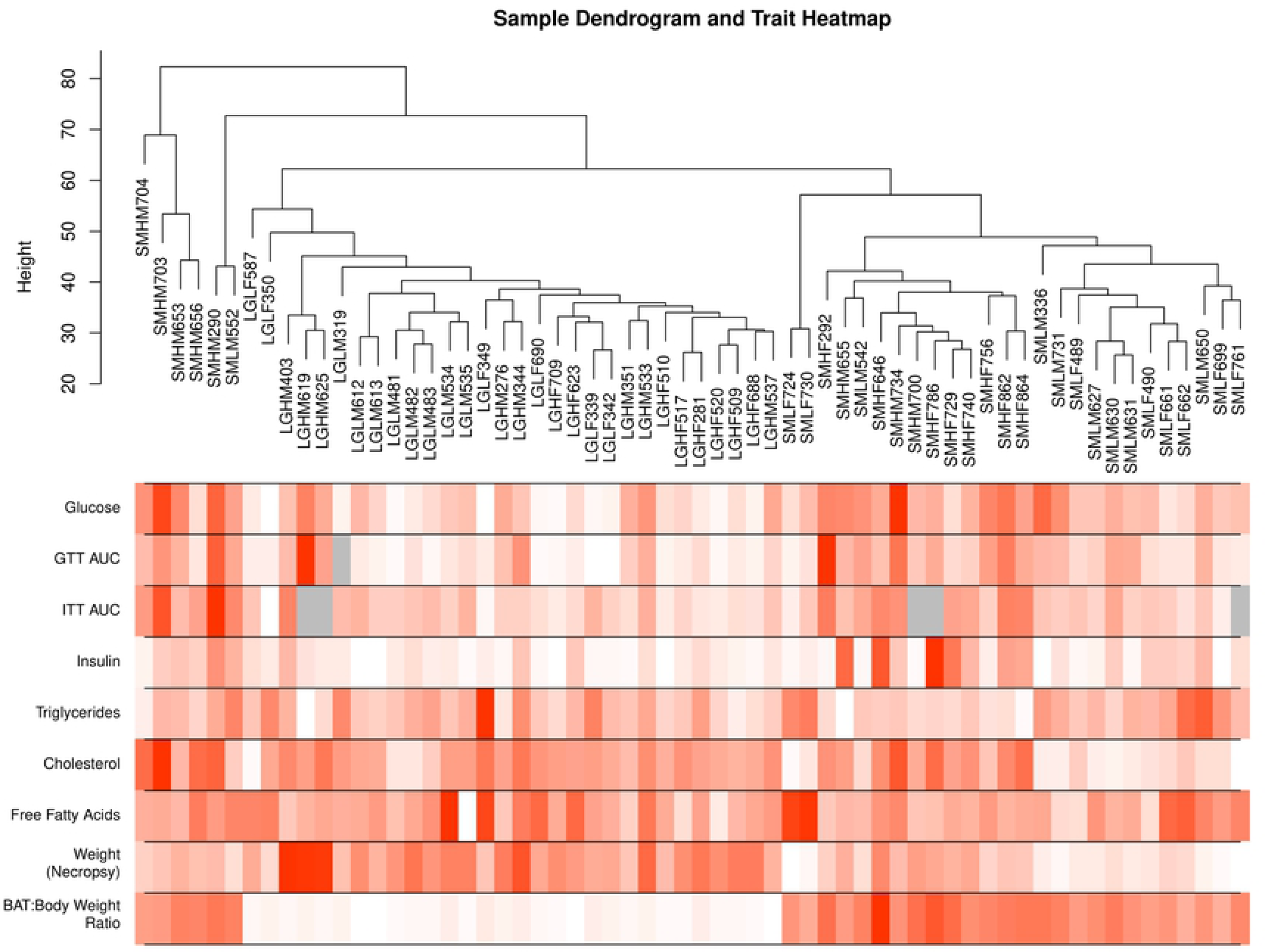
Brown adipose gene expression clustering by strain and diet. Dendrogram of samples clustering based on brown adipose transcriptome (top). All diet by strain cohorts consist of 16 samples (8 male and 8 female) except the low fat LG cohort (8 male and 6 female). Phenotypic values for each sample: red = high value, white = low value, and grey = missing value (bottom). Means for each cohort listed in **Supplemental Table 1**.

### Modules enriched for immune/cytokine response and cell division correlate with glucose parameters and brown adipose to body weight ratio

To identify gene network modules in brown adipose tissue that associate with the observed variation in metabolic phenotypes, we performed WGCNA [53] on the brown adipose gene expression profiles of all cohorts. An unsigned Topological Overlap Matrix assigned genes to 15 discrete modules and eigenvalues were calculated for each module. We then correlated the module eigenvalues with each metabolic phenotype, and find that 8 of 15 modules show significant correlation with at least one phenotype (**Figure 2**). Four modules show significant correlation with at least three of the 4 glucose and insulin parameters (magenta, blue, brown, and pink) and three modules show significant correlations with both body weight and brown adipose to body weight ratio (midnight blue, turquoise and yellow). The blue and brown modules in particular have significant correlations with both glucose phenotypes and brown adipose to body weight ratio.

**Figure 2:**
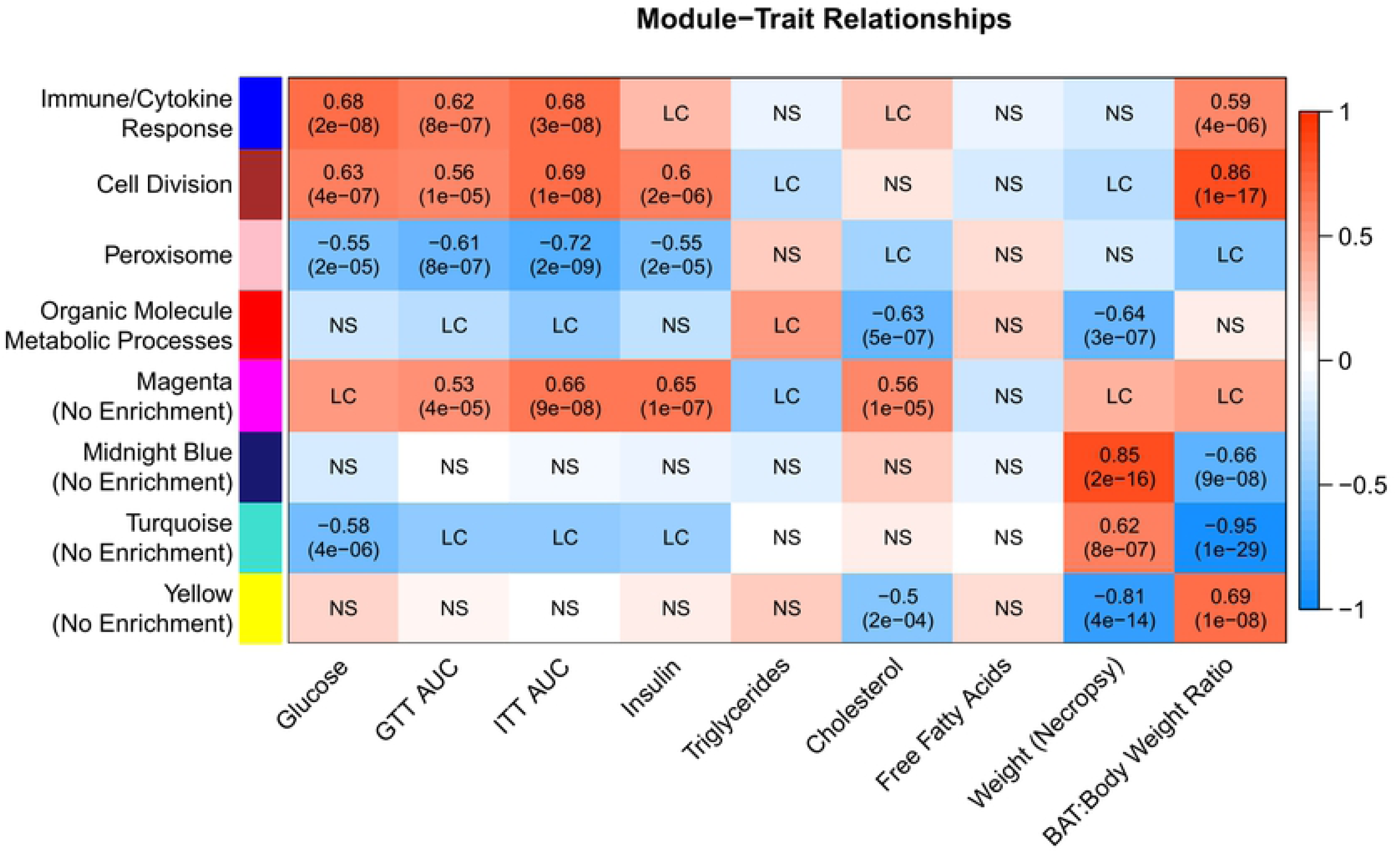
Gene network modules correlate with variation in metabolic traits. Correlation of gene module eigenvalues with metabolic trait values. Enriched GO Terms included in module name. Boxes labeled “NS” showed no significant correlation, while those labeled “LC” had a significant correlation but the strength was between −0.5 and 0.5. Boxes are color coded with strength of the correlation: red = high positive correlation, white = no correlation, blue = high negative correlation.

To determine if the modules are enriched for particular classes of genes, we employed Gene Ontology (GO) term enrichment within each module (**Supplementary Table 2**). Three modules have multiple enriched GO terms and were assigned a general descriptor term for the top 10 enriched terms: blue = immune/cytokine response, brown = cell division and red = organic molecule metabolic processes. The pink module has only one enriched GO term (peroxisome) while the remaining four modules have no significantly enriched GO terms (at an FDR = 0.05). To focus the remainder of our results, we primarily discuss the four modules with enriched GO terms, however all analyses were performed on the modules without enriched GO terms as well.

### Gene expression within network modules varies across both diet and strain contexts

To determine how similar the module-trait relationships are across diets and strains, we analyzed each strain and each diet individually through the WGCNA pipeline (**Supplementary Figure 1, Supplementary Table 3**). At least one immune or immune/cytokine response module is present in both strains and both diets, however the cell division module only appears in the high fat-fed SM/J brown adipose. This led us to hypothesize that some modules may be driven by expression within a particular cohort. To test this, we performed principal components analysis on the genes within each module (**Figure 3, Supplementary Figure 2**).

**Figure 3:**
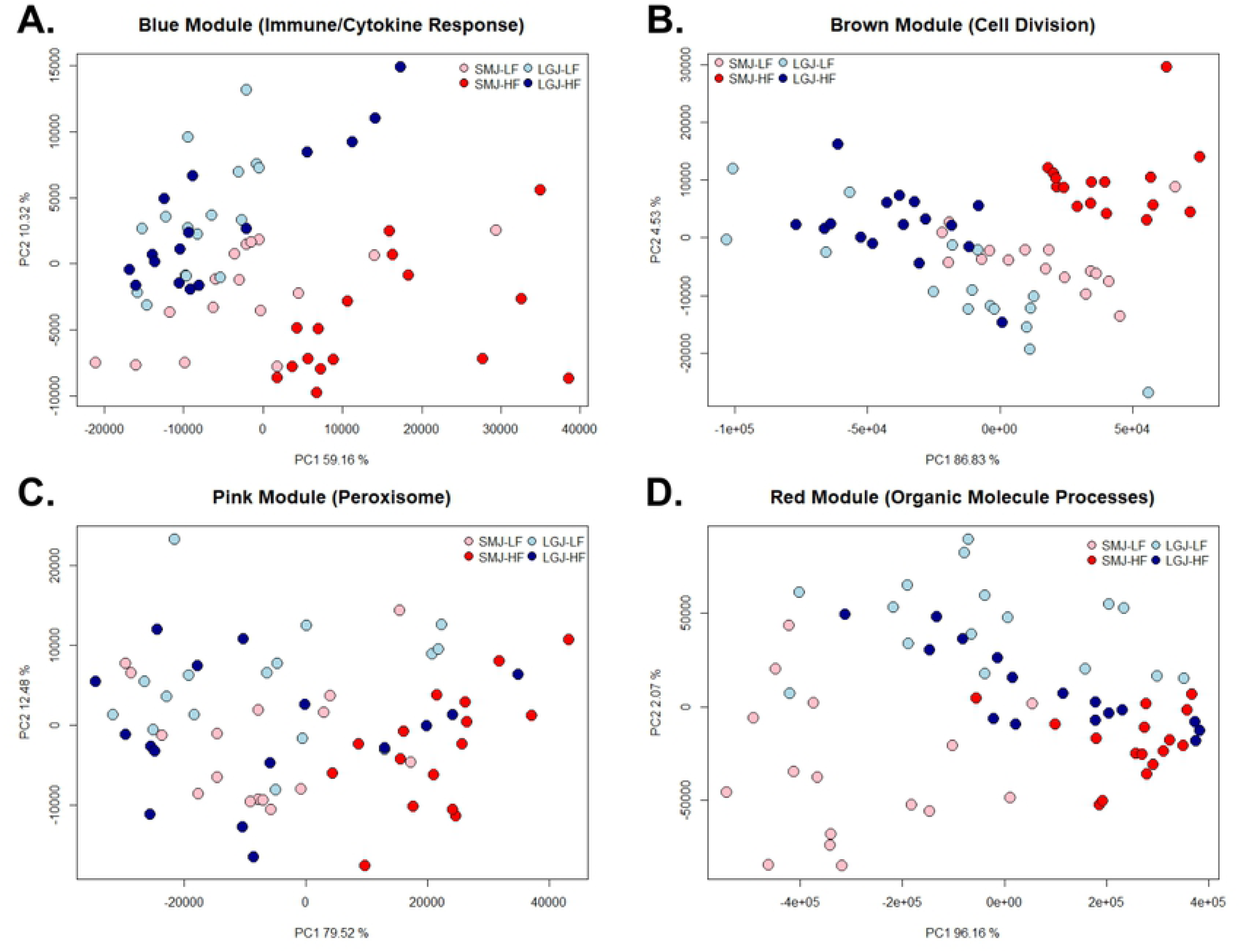
Principal component analysis of each individual module shows variation in diet and strain interactions. Principal component analysis of normalized gene expression counts for each module. (**A**) Variation in the immune/cytokine response module is driven mainly by strain, which further separates into diet. (**B)** High fat-fed SM/J mice stand out in the cell division module. (**C** and **D)** Moderate clustering in the peroxisome and organic molecule processes modules. Samples are color-coded based on strain (LG/J = blue, SM/J = red) and diet (low fat = light, high fat = dark). Plots are labeled with module color and enriched biological process.

Variation in the blue immune/cytokine response module is driven mainly by strain, which then further separates by diet within strain. It is well-established that white adipose tissue is an immunologically active organ that, in obesity, displays both active and adaptive immune responses that affect systemic metabolism [54,55]. However the immunological role of brown adipose tissue is relatively understudied [56]. Our data indicate that genetic background strongly modifies brown adipose tissue’s immunological and cellular signaling processes in response to nutritional environment.

The brown cell division module shows remarkable clustering of the high fat-fed SM/J cohort, which is consistent with our result that enrichment of this term is driven by high fat-fed SM/J mice. Further, high fat-fed SM/J mice have the highest brown adipose to body weight ratios and our previous work showed that the brown adipose expansion observed in these mice is the result of hyperplasia [51]. Both pink peroxisome and red organic molecule processes show moderate diet-by-strain clustering.

### Strain-specific variation drives differential expression and differential connectivity within brown adipose gene modules

To identify the genes that are most differentially expressed between the diets and strains we calculated both differential expression and connectivity. Genes passing an FDR threshold of 0.05 are considered differentially expressed, regardless of fold change. Connectivity was calculated as the degree of co-expression of each gene with all other genes. Genes with differences in connectivity having an absolute value above 0.5 between diets and strains are considered differentially connected. Combining differential expression and differential connectivity between the strains and diets allows us to investigate individual genes in transcriptional networks that are particularly susceptible to differences in diets or genetic backgrounds. To our knowledge this has never been explored in brown adipose tissue.

Genes in the blue immune/cytokine response, brown cell division, and pink peroxisome modules have increased connectivity in high fat-fed mice regardless of genetic background (**Figure 4A**). This indicates that genes in these modules are more tightly co-expressed under nutritional excess. In contrast, the red organic molecule metabolic processes module has a more even distribution of connectivity, with a trend towards higher connectivity in low fat-fed mice. The distribution of differential expression in each module also shows even numbers of genes upregulated in high or low fat-fed mice (**Figure 4A, Supplementary Figure 3A**). However, genes with the highest differential expression tend to also have differential connectivity, with the majority showing increased expression and increased connectivity in animals fed the same diet.

**Figure 4:**
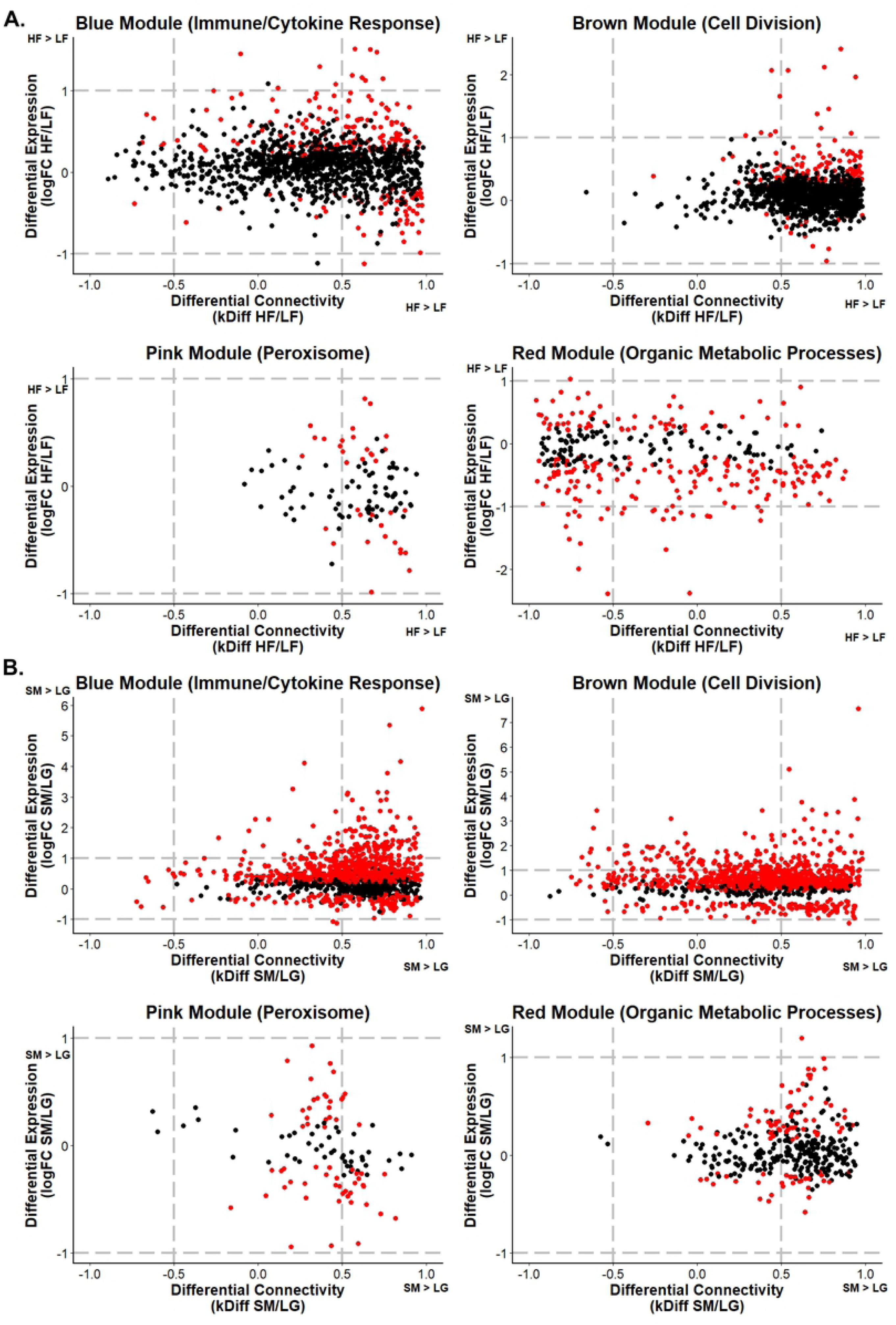
Genes that are both differentially expressed and differentially connected are generally concordant in the direction of difference. Differential expression is plotted along the y-axis as log2 foldchange between (**A**) diets or (**B**) strains (red color indicates FDR-corrected p-value < 0.05). Connectivity is plotted along the x-axis as kdiff between (**A**) diets or (**B**) strains. Genes with a difference in connectivity above 0.5 or below −0.5 were considered differentially connected, and vertical grey dashed lines are provided to visualize these cut-offs. Horizontal grey dashed lines are also provided to visualize log2 foldchange values above 1 and below −1. Positive values indicate higher expression or connectivity in high fat-fed or SM/J cohorts, negative values indicate higher expression or connectivity in low fat-fed or LG/J cohorts.

Analyzing differential expression and connectivity by strain reveals much stronger connectivity in the SM/J strain compared to the LG/J strain in all four of our primary modules (**Figure 4B**) and in two of our four secondary modules (**Supplementary Figure 3B**). Similar to the connectivity-by-diet analysis, genes with high differential expression also have increased connectivity in the same genetic background. However, this may be skewed by the overall quantity of genes that are differentially expressed by strain (n = 4847). To break down connectivity and expression patterns between diets and between strains, we classified genes that showed both differential connectivity and significant differential expression between diets or between strains as potential hub genes. In total, this resulted in 2564 potential hub genes: 659 that are differentially expressed and connected by diet, 2320 by strain, and 415 for diet-by-strain (**Supplementary Table 4**).

To further refine these lists and identify candidates that are likely to be contributing to metabolic variation, we filtered for genes that are differentially connected and differentially expressed with a fold change ≥ 2. This produced a list of 25 genes, 13 of which belong to the blue immune/cytokine or brown cell division modules (**Supplementary Table 5**). Interestingly, these 13 genes are all upregulated and have higher connectivity in the high fat-fed SM/J cohort, indicating that the brown adipose of SM/J mice is particularly responsive to nutritional excess.

This is consistent with our previous work showing brown adipose tissue-dependent resolution of diabetes in high fat-fed SM/J mice [51]. Twenty-four of these hub genes contain small nucleotide variants between the LG/J and SM/J strains [57] (**Supplementary Table 6**), which could be contributing to the gene-by-environmental differential expression patterns we observe. Further, four of these genes, *Col28a1, Bmp8b, Cyp26b1*, and *Kcnj14*, fall within the support intervals of metabolic quantitative trait loci (QTL) mapped in an F_16_ advanced intercross of the SM/J and LG/J strains (**Table 1; Supplementary Figure 4**) [58,59]. These genes represent actionable candidates that can be tested for their function in brown adipose tissue and effects on obesity and systemic metabolism.

## Discussion

Adipose is a dynamic endocrine organ that is critical for regulating systemic metabolism. Further, adipose tissue function displays a high degree of plasticity under different nutritional conditions [60–62]. Though brown adipose has high therapeutic potential for obesity and related metabolic disorders, research on this tissue is in its infancy and most genetic studies focus on identifying the factors involved in its thermogenic function and in determining brown adipocyte identity. Yet recent research reveals that brown adipose is a source of endocrine signals with both anti-diabetic and anti-obesogenic properties [63,64]. High fat diet has been shown to alter brown adipose activity and blunt its positive effects on systemic metabolism [65,66]. Yet, as illustrated by numerous studies, dietary response is heavily dependent on genetic background [16,17,73–75,50,51,67–72]. Here we present the first study on the effects of genetic background and diet on brown adipose transcriptional networks associating with metabolic variation.

We illustrate that the SM/J brown adipose transcriptome is more susceptible to dietary perturbations in comparison with the LG/J strain (**Figure 1**). The LG/J and SM/J strains are frequently used in metabolic studies because they vary in their metabolic response to dietary fat [50,58,59,76,77]. We recently demonstrated that high fat-fed SM/J mice dramatically expand their interscapular brown adipose depots and that contemporary with this expansion, mice enter diabetic remission [51]. Understanding the genetic underpinnings of this phenomenon could open new avenues for understanding novel biology and highlight therapeutic targets for obesity-related metabolic dysfunction.

We identified eight primary gene co-expression modules that are highly correlated with obesity and diabetes traits (**Figure 2**). Four of these modules show significant over-representation of genes belonging to biological categories that affect adipose function and systemic metabolism: immune/cytokine response, peroxisomes, organic metabolic processes, and cell division. Genes involved in immune and cytokine response show a conserved network correlating with glucose and insulin traits (**Figure 3, Supplementary Figure 1**). This is consistent with previous work relating hyperglycemia, hyperinsulinemia, and other diabetes-related traits with inflammatory markers and immune infiltration of adipose [55,78–80]. Peroxisome genes are essential to lipid metabolism, and have been shown to regulate the thermogenic function of both brown and beige adipocytes [81]. Genes composing the organic metabolic processes category include those that perform essential functions in glucose and lipid uptake (**Supplementary Table 3**). Genes involved in cell division form a network specific to high fat-fed SM/J mice, and strongly correlate with glucose and insulin traits as well as with brown adipose to body weight ratio (**Figure 3, Supplementary Figure 1**). The gene-by-environmental specificity of this module, its association with glycemic parameters, and the unique characteristics of brown adipose in the SM/J strain [82], indicate that the genes in this module are compelling candidates for further studies of biological mechanisms of action of brown adipose and systemic metabolism.

We highlight 4 genes that fall in QTL previously mapped in an F_16_ generation of a LG/J x SM/J advanced intercross population (**Table 1**) [58,83]. *Kcnj14*, potassium inwardly rectifying channel subfamily J member 14, is part of the blue immune/cytokine response module. It is an essential membrane protein and inward-rectifier potassium channel that may play a role in glucose uptake in target tissues of insulin. Although *Kcnj14* is understudied, in vivo glucose uptake experiments in mice suggest that disruption of potassium channels affect insulin-stimulated glucose uptake in white adipose [84]. Other studies have shown that potassium channel knock-out mice have hyperlipidemic brown adipose tissue [85]. *Cyp26b1*, cytochrome P450 family 26 subfamily B member 1, is also part of the blue immune/cytokine response module. It is a retinoic acid hydroxylase that regulates cellular concentrations of all-trans-retinoic acid. Retinoic acid is a vitamin A derivative that is essential for cell growth and differentiation. Early studies show that retinoids, including retinoic acid, play an essential role in adipose differentiation [86,87], and a recent study found that retinoic acid mediates adipogenic defects in human white adipose-derived stem cells [88]. *Col28a1*, collagen type XXVIII alpha 1, is part of the brown cell division module. It belongs to a class of collagens involved in extracellular matrix (ECM). The ECM is a critical component in cellular signaling, either through direct interaction with cell-surface receptors or through the ability to regulate growth factor bioavailability [89]. Collagen is highly enriched in adipocytes, and its depletion is associated with metabolic dysfunction [90]. *Bmp8b*, bone morphogenic protein 8b, is part of the red organic metabolic processes module. Of these four candidates, only *Bmp8b* has been studied in brown adipose. It is secreted by brown adipocytes and amplifies the thermogenic response of cells by increasing sensitivity to adrenergic input [91,92].

Gene-by-environment interactions are critical for understanding the intricacies and nuances of obesity and metabolic dysfunction, and for identifying potential therapeutic targets. Though there is increasing interest in brown adipose tissue as a potential therapeutic target for such diseases, there have been no studies on gene-by-environment interactions in brown adipose tissue, and few studies on brown adipose in mouse strains other than C57BL/6J. Our results indicate that gene-by-environment interactions significantly contribute to variation in brown adipose transcriptional networks. Understanding how genetic variation mediates brown adipose tissue’s response to an obesogenic diet will be key to harnessing its therapeutic potential. Further, the unique transcriptomic profile of high fat-fed SM/J brown adipose tissue and its correlation with diabetic remission [51] highlight compelling candidates for understanding brown adipose tissue’s endocrine function and biological mechanisms of action beyond thermogenesis.

## Materials and Methods

### Sample Collection and Sequencing

Experimental animals were generated from SM/J (RRID:IMSR_JAX:000687) and LG/J (RRID:IMSR_JAX:000675) mice obtained from The Jackson Laboratory (Bar Harbor, ME) at the Washington University School of Medicine. All experiments were approved by the Institutional Animal Care and Use Committee in accordance with the National Institutes of Health guidelines for the care and use of laboratory animals. Mice were randomly weaned onto a high fat diet (42% kcal from fat; Teklad TD88137) or an isocaloric low fat diet (15% kcal from fat; Research Diets D12284) and fed *ad libitum*. Additional mouse maintenance details and phenotype collection are described extensively in Carson et al., 2019 [51], and summary statistics for each cohort are provided in **Supplementary Table 1**.

Total RNA was isolated from intrascapular brown adipose tissue using the RNeasy Lipid Tissue Kit (QIAgen). RiboZero (Illumina) libraries were sequenced at 2×100 paired end reads on an Illumina HiSeq 4000. Reads were aligned against LG/J and SM/J custom genomes using STAR [57,93]. Read counts were normalized via upper quartile normalization and a minimum normalized read depth of 10 was required. Additional sequencing and alignment details are provided in Carson et al., 2019 [51].

### Gene co-expression and phenotype associations

The Weighted Gene Co-Expression Network Analysis (WGCNA) R package was used to determine gene co-expression modules and their correlation with metabolic traits [53]. Briefly, edgeR-normalized counts for each gene were converted to a standard normal. Genes with standard deviation of at least 0.25 were deemed biologically variable and used in the subsequent analysis (7740 genes total) [94]. Samples were clustered based on expression of all genes and two low fat LG/J samples were removed as outliers before continuing with the analysis. The adjacency matrix was created from Pearson’s correlations calculated between all genes and raised to a power β of 12, chosen based on a scale-free topology index above 0.9. Raising the absolute value of the correlation by this power is done to emphasize high correlations at the expense of low correlations [95].

The blockwiseModules function was used to create an unsigned Topological Overlap Measure using the adjacency matrix and to identify modules of highly interconnected genes. Each module was assigned a color for identification. Module eigengenes were calculated as the first principal component for each module, and Pearson’s correlations were calculated between each module eigengene and each phenotype to estimate module-trait relationships. Module-trait correlations were considered significant at an FDR-corrected p-value < 0.05 and an absolute correlation of at least 0.5. Gene Ontology term enrichment was calculated for individual modules from a background of all expressed genes in the dataset. Modules were considered enriched for a term at a Bonferoni-corrected p-value less < 0.05. Modules with no significantly enriched terms were designated as “No Enrichment” and modules with multiple enriched terms were classified with an overarching description of the top ten significantly enriched terms.

### Differential Connectivity

Differential connectivity is a measure of the differences in gene interactions between high and low fat-fed mice or between SM/J and LG/J mice. Four subnetworks were created (HF, LF, LG/J, SM/J) with the same 7740 genes as in the full network analysis. Within each subnetwork, we assigned genes to modules and defined module-trait relationships. The connectivity of each gene was calculated in each network with an adjacency matrix as a measure of how correlated the gene is with all other genes in the network. Differential connectivity was calculated between diets or strains for each gene 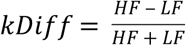 or 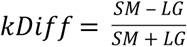 to provide values between −1 and 1. Genes were considered differentially connected when one cohort had three times the connectivity of the other cohort, or an absolute kDiff > 0.5. Genes with positive differential connectivity are more highly connected in the HF or SM cohorts than in the LF or LG cohorts. To further narrow down genes into those most likely to be biologically impactful hub genes, differential expression between diets and strains was calculated for all genes using the exactTest function in edgeR. Genes with an FDR-corrected p-value < 0.05 are considered to be significantly differentially expressed. Hub genes in diet or strain contexts were called as those with both an absolute differential connectivity value > 0.5 and significant differential expression in the same comparison.

## Acknowledgements

This work was supported by the Washington University Department of Genetics, the Diabetes Research Center at Washington University (P30DK020579), the NIH NIDDK (K01 DK095003) to HAL and NIH NIGM (T32 GM007067) to CC.

## List of Supplementary Materials

**Supplementary Figure 1:** Gene network modules correlated with one or more phenotypic trait in individual diet and strain cohorts.

**Supplementary Figure 2:** Principal component analysis of each individual module shows variation in diet and strain interactions.

**Supplementary Figure 3:** Genes that are both differentially expressed and differentially connected are generally concordant in the direction of difference.

**Supplementary Figure 4:** Expression by cohort for four candidate genes

**Supplementary Table 1:** Phenotype summary statistics for each diet-by-strain cohort.

**Supplementary Table 2:** GO term enrichment results for all 8 brown adipose gene network modules.

**Supplementary Table 3:** GO term enrichment results for brown adipose gene network modules in individual diet and strain cohorts.

**Supplementary Table 4:** Differentially expressed and connected genes.

**Supplementary Table 5:** Differentially expressed and connected genes with absolute log fold change ≥1 in both diet and strain comparisons.

**Supplementary Table 6:** List of variants between LG/J and SM/J mice in 24 hub genes.

**RNA sequencing count data available for download at:** http://lawsonlab.wustl.edu/data/

